# The genome of tropically adapted Brahman cattle (*Bos taurus indicus*) reveals novel genome variation in production animals

**DOI:** 10.1101/2022.02.09.479458

**Authors:** Elizabeth M Ross, Loan T Nguyen, Harrison J Lamb, Stephen S Moore, Ben J Hayes

## Abstract

Recent advances in sequencing technology have revolutionised access to large scale genomic data that can be assembled into a platinum quality genome. Here we present a high quality genome assembly with less than 300 gaps of a Brahman cow (*B. taurus indicus*). The assembly was generated using 195GB of PacBio and 169GB of Oxford Nanopore Technologies sequence data. The high quality genome assembly allows us to identify substantial GC content variation that is positively associated with gene rich islands, and negatively associated with genetic variation in the form of structural variants. In addition, 92371 structural variants that are segregating in the brahman population were identified. Gene ontology analysis revealed that genes with varying copy numbers were enriched for gene ontology terms related to immune function. This analysis has revealed the complex structure of the mammalian genome of an outbred species, and identifies the ability of long read data from diploid species can be used to not only assemble a high quality genome, but also discover novel genetic variation within that genome.

## Introduction

Commercial cattle consist of two sub species, the European style *Bos taurus taurus*, and the Asian style *B. taurus indicus*. While *B. taurus* breeds are popular in milk production systems (e.g. Holsteins) and many beef production systems (e.g. Angus), they perform poorly in tropical climates due to heat stress and significant ecto-parasite burdens. In tropical environments *B. indicus* cattle outperform *B. taurus* cattle. One breed, Brahman, are used extensively for meat production in Australia and central and south America. Brahman cattle experience low tick burdens [1] and lower heat stress that *B. taurus* breeds [2, 3], allowing them to thrive in hot, humid and otherwise challenging environments.

Studies on Brahman cattle have historically necessitated the coercion of the genetic structure of the breed into a *B. taurus* framework,with the current standard genome assembly being from a *B. taurus* cow [4]. This causes a number of issues in genomic studies such as difficulty accurately identifying and locating genetic variants which in turn is needed to characterise quantitative trait loci (QTL). Any differences between the true SNP location in Brahman and the assigned location in the *B. taurus* reference will cause additional noise in the results, which translates to wider QTL peaks and an inability to narrow down lists of candidate genes.

The price of sequencing has been decreasing exponentially since high throughput sequencing became popular. While next generation sequencing is broadly used to describe any sequencing that is high throughput – which is generally all sequencing except Sanger sequencing – recently long read high throughput sequencing has become more popular. The two main contributors to long read high throughput sequencing are Oxford Nanopore Technologies (ONT) and Pacific Biosciences (PacBio). These technologies routinely result in reads greater than 10,000bp long. However this long length is at a cost of accuracy in comparison to short read Illumina data. Favourably, ONT and PacBio data accuracy does not appear to decrease with longer read lengths (unlike Illumina), and the errors within PacBio date does not have a specific bias [5]. This means that, theoretically, with enough sequence depth and long reads the true underlying genome sequence can be assembled in its entirety.

Here we aimed to generate a breed specific high quality genome assembly for Brahman cattle using both PacBio and Oxford Nanopore long read data. Using this genome assembly we are able to examine genome structure in previously unachievable detail.

## Results

### Assembling a highly continuous genome

A total of 195Gbp of sequence data was obtained from the PacBio Sequel. The N50 of an assembly is the contig (or reads) length at which 50% of the data is contained within contigs (or reads) at least that long. The mean polymerase N50 was 20.5 kbp ± 0.3kbp across 28 SMRT cells, the mean subread N50 was 17.3 kbp ±0.12. After scrubbing with the DAZZler (DAZZler https://dazzlerblog.wordpress.com/) scrubber pipeline and assembly with FALCON, 1867 primary contigs were obtained. The total length of the assembly was 2,673,704,821bp (Table 1). The contig NG50 for the raw assembly was N50 was 11,122,567bp. The longest contig was 56,814,201bp; the longest 69 contigs account for half of the genome length (LG50); the 20 longest contigs account for 24.6% (689,239,016bp) of the total genome length (25.7% of the assembly length).

**Table 1.**
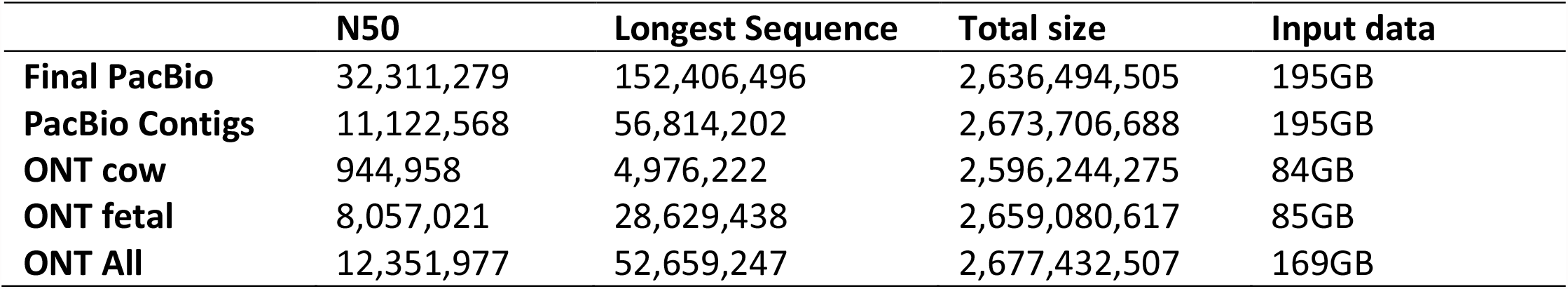
Genome assembly metrics for the Brahman cow using PacBio and ONT sequencing data. ONT assemblies were post-process with RagTag [8] to break inter-chromosomal chimeras.

|  | **N50** | **Longest Sequence** | **Total size** | **Input data** |
| --- | --- | --- | --- | --- |
| **Final PacBio** | 32,311,279 | 152,406,496 | 2,636,494,505 | 195GB |
| **PacBio Contigs** | 11,122,568 | 56,814,202 | 2,673,706,688 | 195GB |
| **ONT cow** | 944,958 | 4,976,222 | 2,596,244,275 | 84GB |
| **ONT fetal** | 8,057,021 | 28,629,438 | 2,659,080,617 | 85GB |
| **ONT All** | 12,351,977 | 52,659,247 | 2,677,432,507 | 169GB |

Contigs were scaffolded with 43GB of Chicago data and 41GB of Hi-C (Dovetail Pty Ltd). Both Hi-C and Chicago libraries were made from DNA extracted from the reference animal, and sequenced on a 2×151bp Illumina platform. The HiRise software (Dovetail Genomics) made 850 and 256 joins from the Chicago and Hi-C data respectively. Mapping these scaffolds to the *B. taurus [4]* genome chromosomes; each chromosome was present in between 1 and 3 ultra-long scaffolds. A subsequent round of scaffolding with the Hi-C data was able to assemble each chromosomes into a single scaffold.

Chromosome scaffolds were polished with the original PacBio sequence reads then gaps were filled using the same reads by PBJelly [6]. A total of 5 rounds of polishing and gap filling was performed before no additional gaps were filled. ONT data from a Brahman bull was aligned to the assembly and assembly regions flanking gaps were examined. Three gaps were completely spanned by reads, the first was a 8.6kb repeat on chromosome 1, the second was a 5.5kb insert, the third showed an incorrectly inserted 7.7kb region that was replaced with 320bp.

After gap filling 317 gaps remained on the assembled chromosomes (with 86 in unplaced scaffolds). Inspection of the 50kb of sequence either side of the remaining gaps revealed that the gaps tended to be surrounded by highly repetitive or duplicated sequences. This may indicate that the inability to fill the remaining gaps is due to complex regions of the genome, and the extremely long sequence reads may be required to close the remaining gaps, as the reads must span long repetitive stretches.

DNA from a liver sample of the same animal was sequenced on the Oxford Nanopore Technologies MinION and PromethION and produced 84GB of sequence data (approximately 31X coverage). Additionally, DNA from liver and lung of the foetal daughter of the Brahman cow animal was also sequenced on the ONT MinION and PromethION. An additional 85GB of data was generated. This data was assembled with Raven [7] and generated 2.6GB of contigs (Table 1). The assembly of the cow data had a shorted N50 than the foetal data, which had a longer read N50. When the ONT data from the cow and foetus were assembled together, a higher N50 than either one alone was achieved.

### GC content varies between and within chromosomes

An important difference between long read genome assemblies and short read genome assemblies is the GC bias in the raw data. Along with the ability of long reads to avoid the collapsing down of repetitive elements, this provides an opportunity to examine the distribution of GC across the genome. The GC content of each 1MB non-overlapping window was calculated using EMBOSS GeeCee [9]. The GC content of the blocks ranged from 36-60% (Figure 1) in a non-normal distribution (Shapiro-Wilk test, p<0.0001), with a mean and median of 41% and 41.8% respectively. Additionally, there were significant differences between the GC content of chromosomes (Kruskal Wallis test; p < 0.0001), with chromosomes 19 and 25 having the highest median GC content (46%), and chromosomes 1, 6 and 9 having the lowest median GC content (39%).

**Figure 1.**
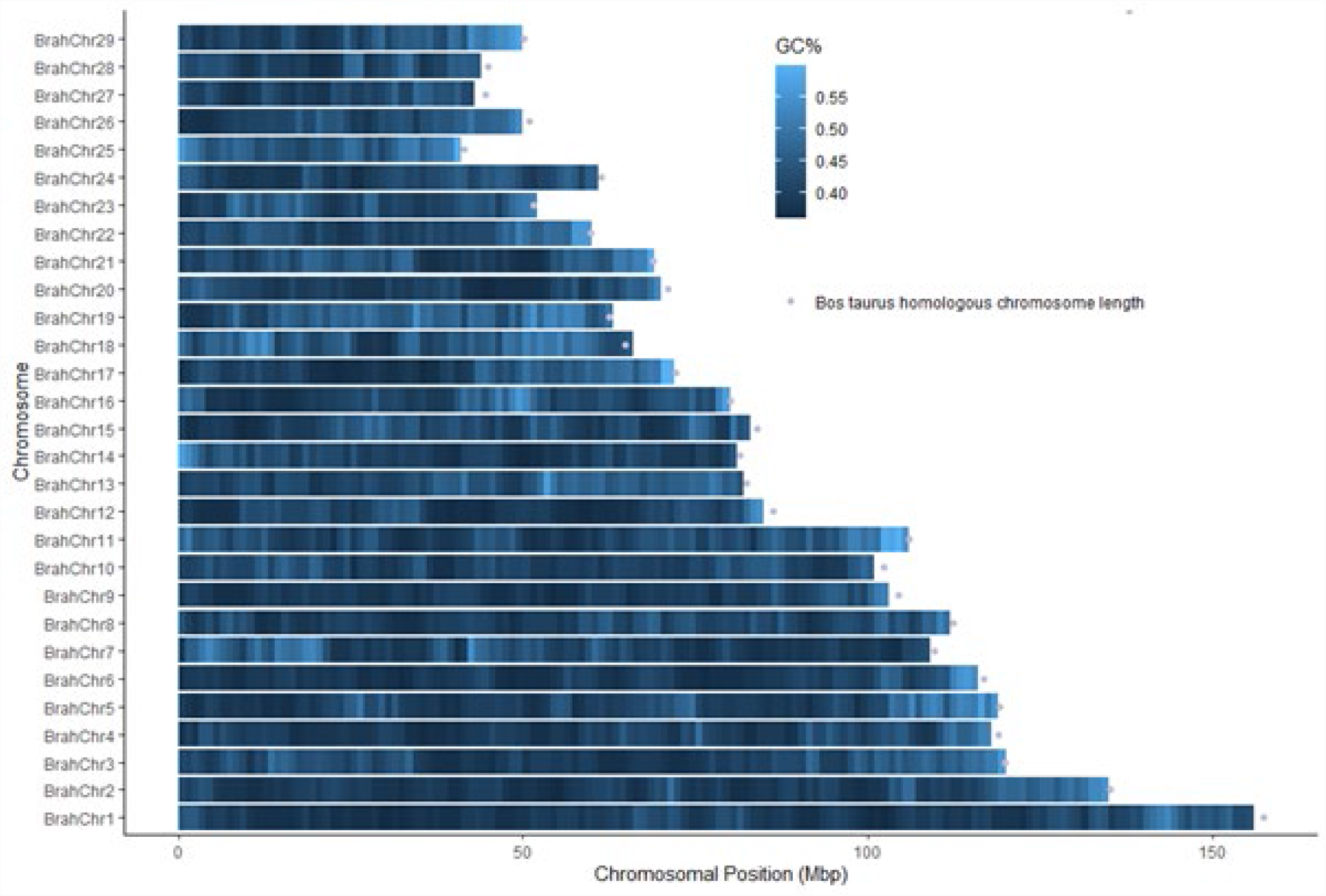
GC% of 1MB non-overlapping windows. *B. taurus* chromosomes lengths taken from ARS 1.2 [4]. GC of each window was calculated using GeeCee [9].

### Structural variation in the Brahman genome

In total 92371 structural variants of more than 10 base pairs were identified, accounting for 0.973Mbp of sequence (Table 2; Figure 2). Almost all (>99.9%) of the identified structural variants were classified as insertions (50.7%) or deletions (49.3%). Of the identified insertion and deletions 77.0% and 72.8% were less than 50bp in length. Only 4.6% of all structural variants were longer than 500bp, and all of these were insertions or deletions. For small indels that were less than 50bp long there were more insertions than deletions (p<0.0001), for intermediate indels there where equal numbers of insertions and deletions (p>0.05), and for long indels of over 5000bp there were only deletions. Very small indels – those less than 20bp in length, accounted for just over half of all the identified variants (50.5%), but only made up 6.26% of the total base pairs contained in structural variants. While smaller structural variants were more common, larger structural variants, above 500bp, accounted for more than half of the base pair located within structural variants (56.3%).

**Table 2.** Size of structural varients observed within the heterozygous Brahman genome

| **SV Size** | **Insertions** | **Deletions** | **Other** |
| --- | --- | --- | --- |
| **>10 bp** | 46849 | 45508 | 14 |
| **>50 bp** | 10774 | 12393 | 8 |
| **>500 bp** | 2769 | 1518 | 0 |
| **>5000 bp** | 0 | 66 | 0 |

**Figure 2.**
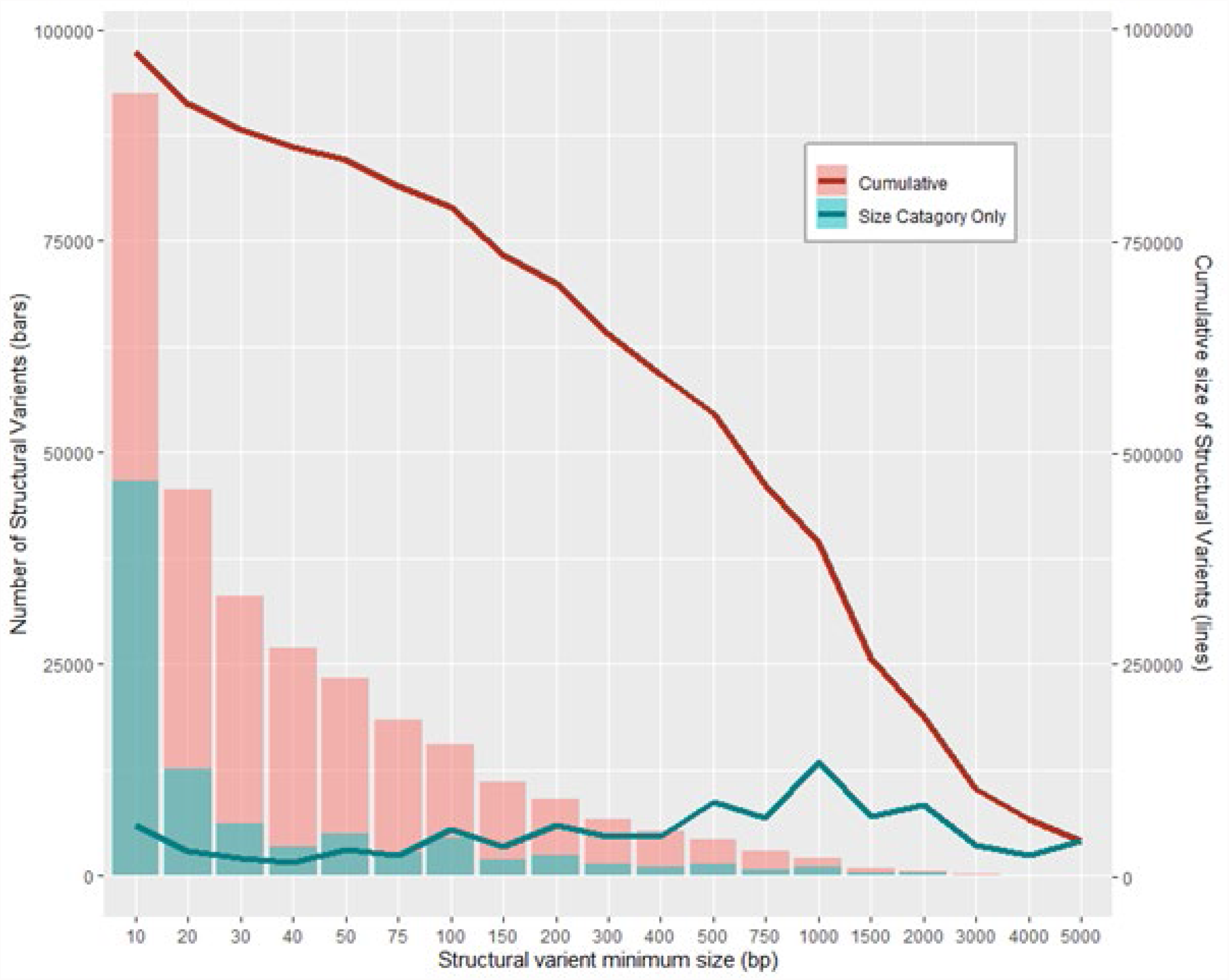
Structural variants in the Brahman genome.

### Genome regions are enriched for genes and functions

Genes from the EMBL annotation of the *B. taurus* genome were located in the Brahman genome using BLASTn. To be considered correctly placed within the Brahman genome the genes had to have a single location and an identity between the source *B. taurus* genome and target Brahman genome of at least 97%. Of the annotated genes, 14845 had a significant hit to the Brahman genome and 14219 (95.8%) genes were uniquely placed, 526 (3.5%) had between two and ten significant hits, and 100 (0.7%) had significant hits to more than ten locations within the genome. Of the uniquely placed genes 90% had an identity of at least 99% (Table 3). A GO term enrichment analysis indicated that genes that map to more than one location were enriched for 474 functions consistent with transposable elements such as “DNA recombination” and “RNA polymerase II general transcription initiation factor activity”. Genes which were uniquely placed in one location of the Brahman genome shared a higher proportion of similarity between the Brahman and taurine genome than the genes which were non-uniquely placed (p<0.001).

**Table 3.** Gene ontology enrichment of *B. taurus* genes placed within the brahman genome.

|  | **Genes** |  | **Gene Ontology<sup>1</sup>** |  |  |
| --- | --- | --- | --- | --- | --- |
|  | Count | % of unique | CC | BP | MF |
| **Uniquely placed** | 14219 | 100% | 1150 | 9045 | 1754 |
| **Not uniquely placed** | 84192 | 0% | 101 | 270 | 103 |
| **Identity ≥ 99.9%** | 1428 | 10.04% | 5 | 63 | 21 |
| **Identity < 99.9%** | 12791 | 89.96% | 10 | 14 | 11 |
| **Identity ≥ 99%** | 12842 | 90.32% | 38 | 111 | 46 |
| **Identity < 99%** | 1377 | 9.68% | 14 | 69 | 45 |
| **Identity ≥ 98%** | 14059 | 98.87% | 2 | 5 | 5 |
| **Identity < 98%** | 160 | 1.13% | 11 | 25 | 13 |
| **Identity ≥ 97.5%** | 14170 | 99.66% | 1 | 2 | 3 |
| **Identity < 97.5%** | 49 | 0.34% | 1 | 2 | 5 |
| **Genes > 20Kbp** | 4074 | 28.65% | 170 | 448 | 174 |
| **Genes > 100Kbp** | 350 | 2.46% | 57 | 156 | 68 |

<sup>1</sup>To calculate Gene ontology significance of unique/non-uniquely placed genes the entire gene set was used; to calculate Gene ontology significance of homology levels and gene length only the uniquely placed gene set was used. BP = Biological Process; CC = Cellular Component; MF = Molecular Function.

The average density of uniquely placed genes varied between chromosomes, with chromosomes 18,19 and 25 the most gene rich. Within each chromosome significant clusters of genes were observed on all chromosomes (p<0.0001) (Figure 3). The MB windows with significantly more genes than expected by chance (> 99.9th percentile of a random distribution based on number of genes and length of each chromosome) also had significantly higher GC values than the other windows (t-test, p< 0.0000001).

**Figure 3.**
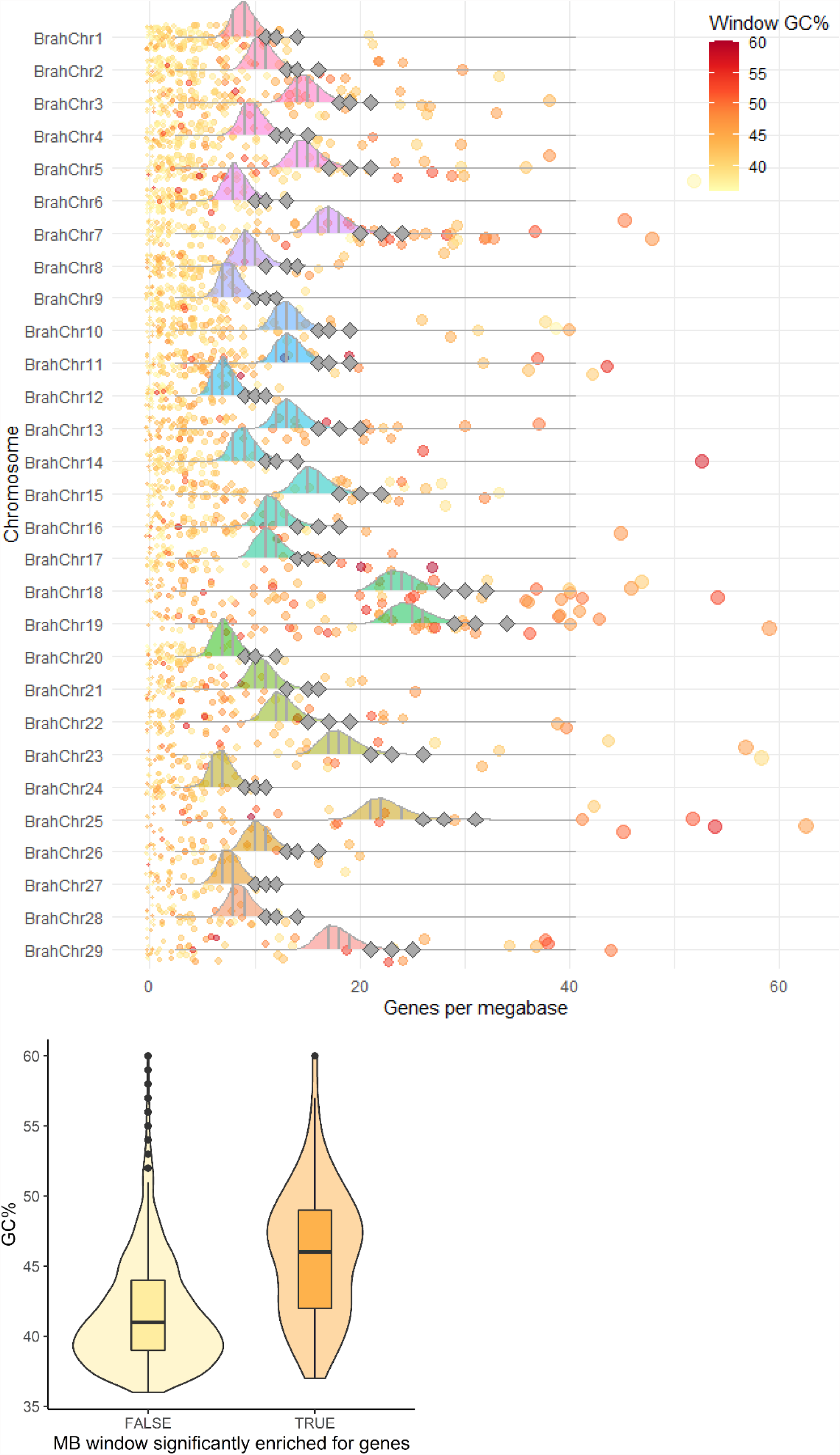
Gene islands in the Brahman genome. A) Distribution of observed uniquely placed genes in each 1MB (round dots) window compared to the distribution expected if genes are randomly dispersed along the chromosome. The 95, 99 and 99.9 percentiles from the random distributions for each chromosome are shown as grey diamonds. B) GC content of windows that have gene content greater than the 99.9th percentile of the random distribution.

### High GC regions have fewer structural variants and more genes

There were significantly less structural variants in the 1MB windows with GC > 40% than those with GC% ≤ 40% (p = 0.0052; Figure 4). The opposite pattern was observed for uniquely placed genes, with more genes observed in the 1MB windows with GC% > 40% (p = 2.2 × 10^−16^); however there were again less non-uniquely placed genes in higher GC% windows (p = 6.6 × 10^−16^). There was a significant negative relationship between the number of unique and non-unique genes in a 1MB block (Kendall’s rank correlation test tau=-0.116 p=2.815 × 10^−15^; Figure 4), a significant negative relationship between the number of uniquely placed genes and structural variants (Kendall’s rank correlation test tau=-0.1778 p=2.2 × 10^−16^), and a significant positive relationship between the number of non-uniquely placed genes and structural variants (Kendall’s rank correlation test tau=0.06221 p=1.109 × 10^−5^).

**Figure 4.**
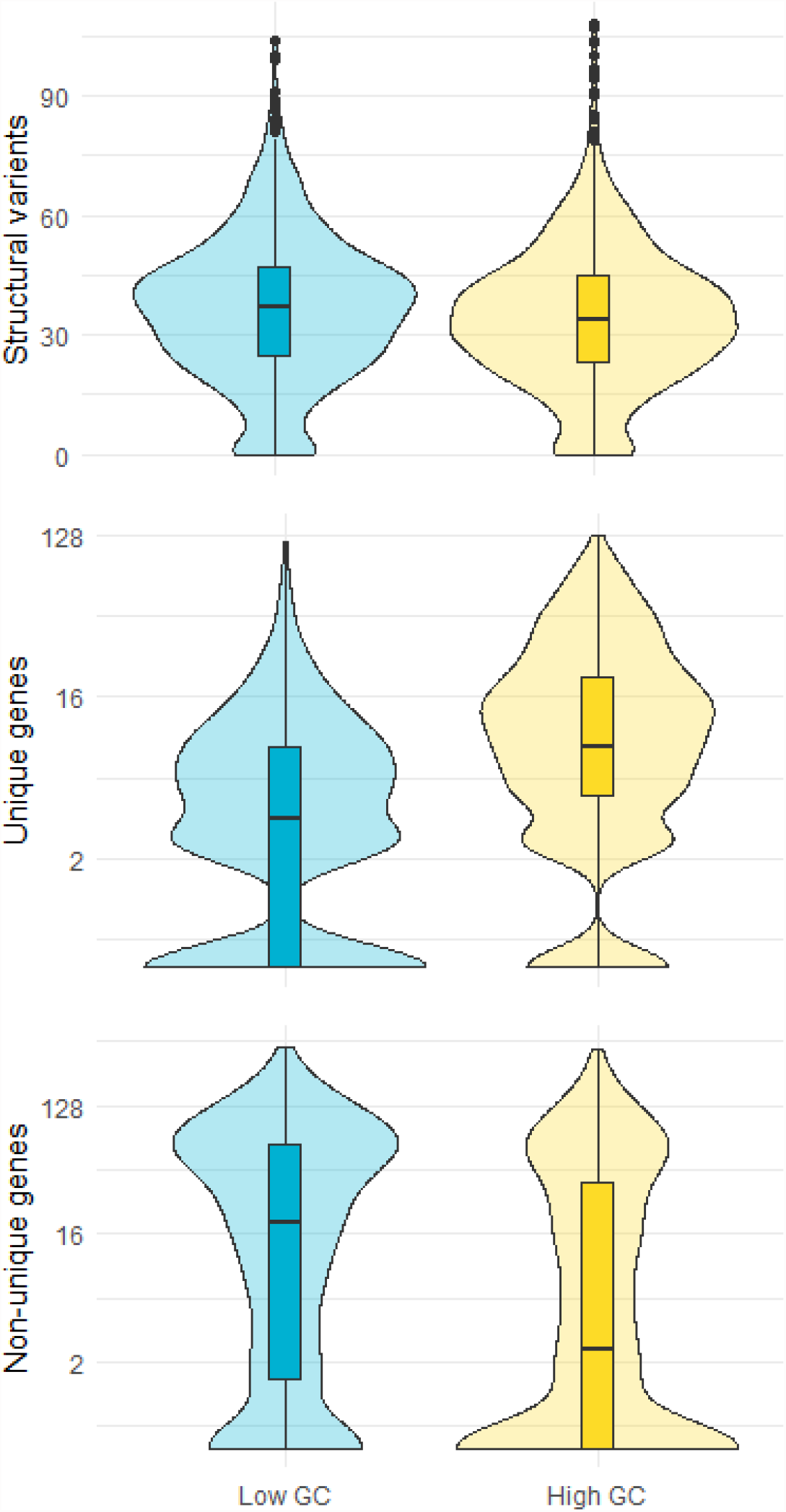
Distribution of the number of a) structural variants, b) Uniquely placed genes and c) non-uniquely placed genomes in high (GC>40%) and low (GC<40%) 1MB regions of the Brahman genome.

### Divergent and conservative selection in gene function

Within the 14219 genes which were uniquely placed within the Brahman genome, 90% had an identity of at least 99% (Table 3). Only 75 genes did not have any homology to the Brahman genome based on the BLASTn alignments, and 75 were placed within the genome but with a homology level lower than the 97% cuttoff. Only 1.1% of uniquely placed genes had an identity of 98% or less. The two extremes of <98% and >99.9% were considered to be conserved and divergent genes.

A total of 84 gene ontology (GO) terms were significantly associated with conserved genes (homology of over 99.9%) (Table 3; Figure 5A), and 49 were associated with divergent genes (homology <=98%). Two GO terms, GO:0004984 and GO:0050911 were enriched in both conserved and divergent gene sets (Figure 5B). These two GO terms share the majority of assigned genes (Figure 5C), and belong to functions related to olfactory sensing and detection of chemical stimulus.

**Figure 5.**
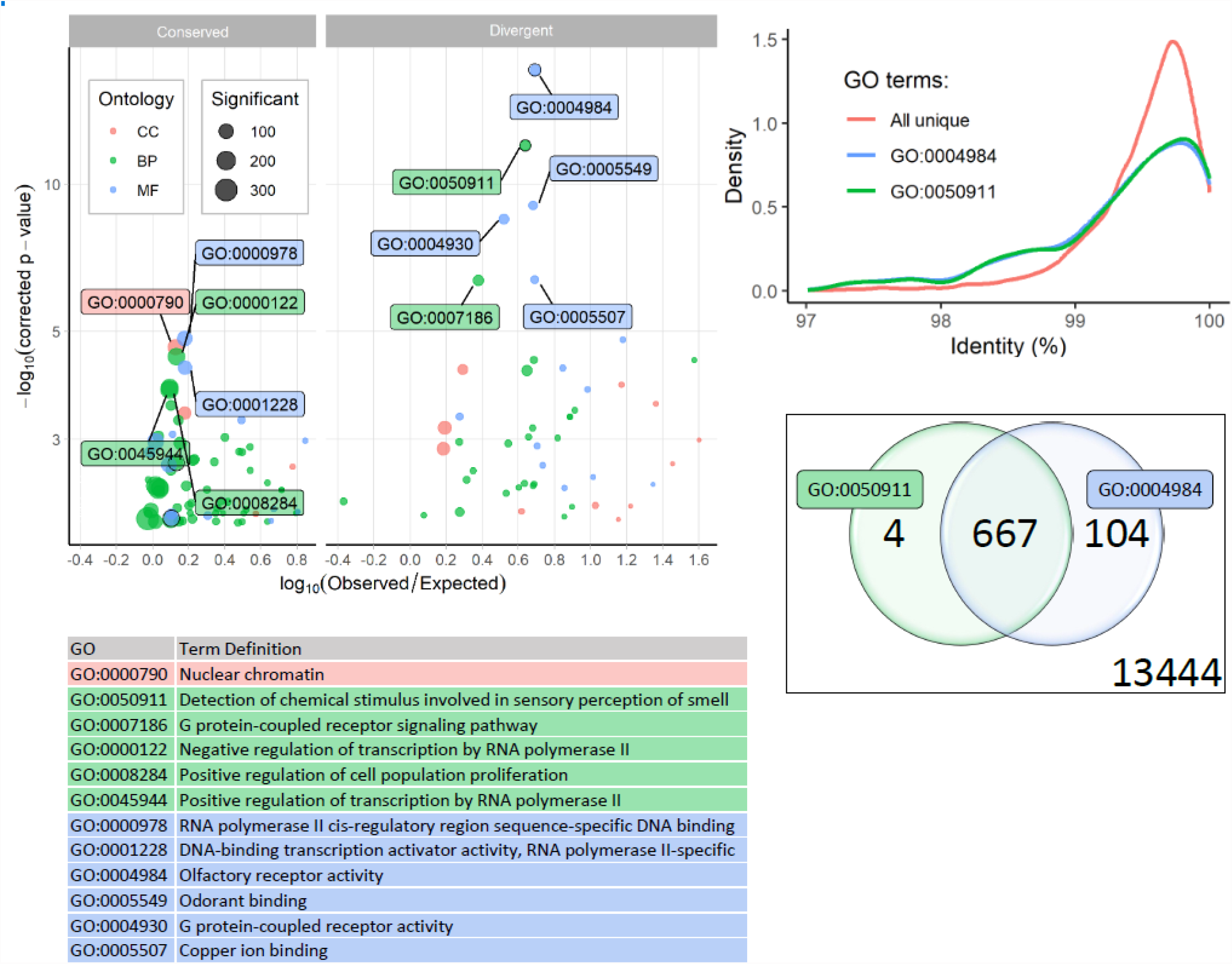
Gene Ontology (GO) enrichment of genes which are conserved and divergent between the *B. taurus* and the Brahman genome. A) Top GO terms enriched in the Conserved and divergent gene sets. B) Definitions for top GO terms in the Conserved and divergent gene sets. C) Distribution of the identity between the two GO terms that are enriched in both conserved and divergent gene sets. D) Overlap of genes in the two GO terms that are enriched in both conserved and divergent gene sets. BP = Biological Process; CC = Cellular Component; MF = Molecular Function.

### Duplicated genes in the Brahman genome

To identify genes that have been duplicated in the Brahman genome compared to the Taurine genome, the number of significant BLAST hits of each gene was compared between the two assemblies. There were 46 genes with a higher number of significant BLAST hits to the Brahman genome than the taurine genome; there were 53 genes with a significant number of BLAST hits higher by two or more in the taurine compared to the Brahman genome. Within these genes, there were 145 and 105 enriched GO annotations compared to the whole genome for the genes with more hits to the Brahman and taurine genomes respectively (Figure 6). In the genes which had higher copy numbers in the Brahman genome the GO terms with the most significant p-values were olfactory receptor activity and G protein-coupled receptor activity. Additionally a number of immune response related terms were present, including adaptive immune response, regulation of immune response, response to virus, “natural killer cell activation involved …”, neutrophil degranulation, defense response to virus, antibacterial humoral response, B cell proliferation, “regulation of defense response to virus …”, T cell receptor complex, “T cell activation involved in immune res…”, B cell differentiation and humoral immune response. In the genes with higher copy numbers in the taurine compared to the Brahman genome enriched GO terms included positive regulation of RNA interference and vitamin D receptor binding. There were no GO terms in common in the enriched lists between the expanded and contracted gene lists.

**Figure 6.**
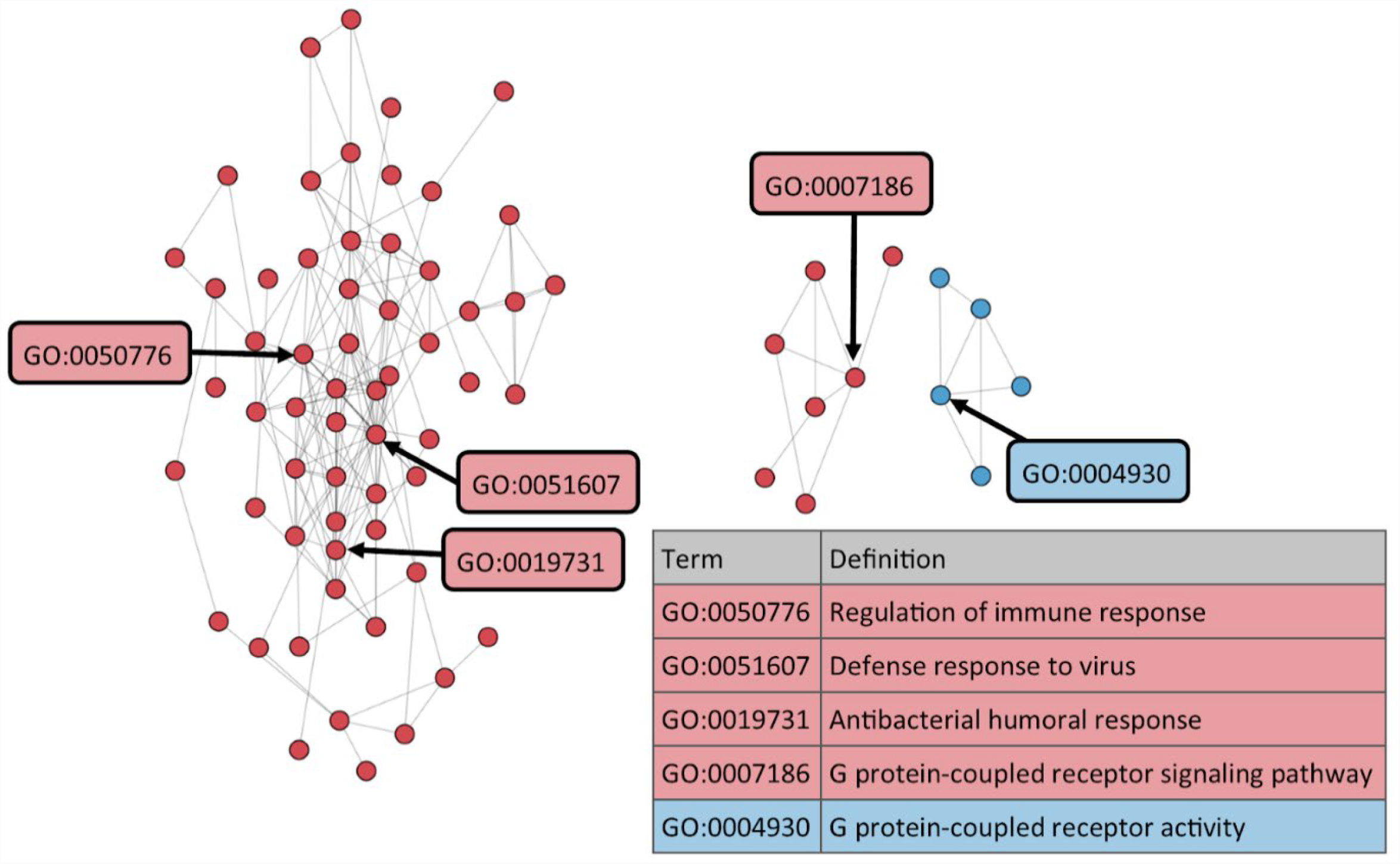
Functional network of GO terms significantly associated with genes that have different copy numbers between the Brahman and taurine genomes. Only clusters with more than 4 nodes are shown. Distance measure: RSS. Cutoff 0.6. Relationships were generated using NaviGO using a list of GO terms identified as significantly enriched in the expanded and reduced gene list.Table 1. Genome assembly metrics for the Brahman cow using PacBio and ONT sequencing data. ONT assemblies were post-process with RagTag [8] to break inter-chromosomal chimeras.

A network analysis of the duplicated genes were undertaken using NaviGO. Three clusters of functions were identified. The largest cluster contained central nodes with functions related to immune activity, while the other two smaller nodes related to G-protein-coupled receptor activity.

## Discussion

Here we present the diploid genome of the tropically adapted Brahman (*B. taurus indicus*). The genome revealed that there is extensive variation in GC content throughout the genome, and that this variation in GC content is associated with the abundance of structural variants and genes. Significant functional enrichment was found for pathways involved in response to chemosensory stimuli - in both conserved and divergent genes. Novel gene duplication events were identified, and genes with differing copy numbers between the taurine and Brahman genomes were enriched for functions related to immune response.

Here we have used two long read sequencing methods to assemble the genome of the same animal. Both methods were able to produce contigs in excess of a megabase in length. Comparison between the PacBio and ONT assemblies revealed a number of inter-chromosomal chimeras, that were confirmed by mapping contigs to the PacBio assembly and the *B. taurus* assembly. These chimeras appeared to be relatively rare, and could be easily resolved using a program such as RagTag [8]. This suggests that assembling a genome with ONT data using Raven should be carefully examined, and any putative chromosomal rearrangements should be interpreted with skepticism. However, the concurrence within each chromosome between the two methods was high, providing a good avenue for cost effective breed specific genome assemblies that will allow the examination of structural variants and gene duplications, at which short read sequencing methods are notoriously poor.

Even in the most well studies genomes some regions remain elusive. For example, the human genome assembly has 651 unfilled gaps [10]. Here we used two independent datasets generated from the same individual, and related individuals, to minimise gaps within the assembly. This approach was highly successful, however interestingly some regions showed gaps in genome assemblies in from both technologies. This suggests that rather than gaps in genome assemblies being due to random regions of low coverage, that rather there are segments of the genome that are particularly difficult to assemble even with modern long read technologies. Potentially this phenomenon could be caused by long repetitive stretches, where the reads cannot span the repeat region.

We have presented 92371 structural variants that are segregating in the Australian Brahman population. These structural variants account for an entire megabase of variable bases within this one animal. Structural variants are known to cause some dramatic phenotypic effects, including the lack of horns in and coat color cattle, and a range of phenotypic associations in humans and model organisms. Currently structural variants are not used extensively for phenotypic prediction of quantitative traits in livestock, however the ability to characterise and genotype known structural variants may mean that the phenotypic variation that they induce can be captured by genotyping them using imputation from SNP arrays in the future. Long read data in cattle has already been used to identify economically important traits in cattle such as the poll (hornless) mutation [11], coat color patterns (under review), and deleterious mutations [12].

GC variation across the Brahman genome has been characterised here. The ability to characterised genome wide variation in GC content is a direct consequence of the sequencing technology used. Short read Illumina sequencing, which was used extensively to build the first generation of livestock reference genomes has a known GC bias, which makes it impractical for identifying GC rich and poor regions [5]. PacBio and ONT data do not share the same bias, and so we were able to identity significant chromosomal variation in GC content, as well as substantial within chromosome GC variation. Sections of high and low GC content is a known occurrence across many eukaryotic species [13], however early reports of interchromosomal total GC variation in humans [14] likely underestimate the amount of variation due to the bias in short read technology.

There is a significant relationship between the GC content of a 1MB window and the number of genes it contains. Additionally we noted that there are less structural variants in regions with higher GC content. Whether this is due to active selection associated with maintaining gene function, and thus limiting the introduction which may interrupt the correct function of genes, or whether there is a relationship between the introduction of a structural variant and GC contents itself is currently unknown.

Genes were found to be clustered within the genome in a non-random fashion, often clustering in gene rich islands. Eight gene rich islands that had over 50 genes within a 1MB window were identified. These gene rich islands were also found to have a significantly higher GC content. This may be an effect of tandem duplication leading the novel genes, but it also may be related to co-expression of genes within a pathway, whereby open chromatin within a region can allow the expression of a range of genes which are required for a certain function. The GC content in a region has been shown to be associated with the physical loop topological associated domains (TADs) in mammalian cells [15]. In turn, TADs have a role in the way gene regulation is controlled [16], suggesting that these regions may be co-expressed.

Due to the number of genes within the gene rich islands, it is possible that these regions may have a higher than random effect of the phenotype of the animals. For animal breeding and gene association based studies additional SNP markers may be included in SNP arrays within these gene rich islands, to ensure practices such as imputation (where full sequence markers are inferred from lower density markers based on linkage disequilibrium), can achieve high levels of accuracy in these regions. While the idea situation for causal gene identification is the have a marker on the array in perfect linkage disequilibrium with the causal variant, a more likely scenario is that the causal marker is imputed from low density information. Importantly, imputation errors can lead to reduced power in genomic association studies [17] due to the introduction for additional errors (noise), and thus may lead to identification of incorrect causal variants. Marker density has a known effect on imputation error [18], and correcting incorrectly placed markers may further reduce error [17]. Hence, care should be taken to curate markers within these potentially phenotypically important domains.

Genes in the Brahman genome were identified based on homology to the *B. taurus* gene set. Most (90%) genes had high homology, >99%, which is concurrent with the relatively recent evolutionary split between *B. indicus* and *B. taurus*. Genes for olfactory signalling were enriched in both the divergence and conserved gene sets, suggesting there may be selection pressure acting on these pathways. Koufariotis *et al*. [19] found that genes for olfactory detection were enriched in genome regions of brahman cattle that had significant *Bos taurus* introgression, and studies in other species have found that olfactory related genes undergo rapid evolution [20, 21]. Taken together, this pattern could be explained by olfactory receptors rapid evolution, but taurine versions being interogressed in some genome regions giving the effect of both highly divergent and conserved (because they are introgressed) homology.

Genes which had evidence of duplication within the Brahman genome compared to the taurine genome were enriched for immune functions. Gene duplications are thought to be a mechanism for adaptation [22]. Given the significantly different ability of the two subspecies to mount a successful immune response to ectoparasite burden [1, 23], the subspecies specific gene duplications may be reflective of the difference biotic stresses that each has adapted to.

This study has examined the structure of an outbreeding mammal. As genome assemblies become more economical to produce it will be possible to validate these findings across the mammalian kingdom. However, the use of multiple datasets here has demonstrated the danger of assuming an assembly is correct, as seen by the inter-chromosomal chimera produced by the raven assembly. Best practise should use multiple animals or data types to confirm any large scale variation in genome structure, especially as datasets become easier to produce.

This analysis had revealed the structural patterns within a cattle genome, and demonstrated the genetic insights that can be found by generating high quality breed specific reference genome assemblies. As the costs associated with these technologies continue to decline it is foreseeable that each economically important livestock breed will have a tailored genome assembly, increasing the development of genomic tools and technologies to advance genetic improvement in livestock and increase our understanding of the mammalian genome and its function. Here we present an overview of patterns, however comparative genomics of this and other genomes are likely to further illuminate the complexities of the mammalian genome.

## Methods

### Animal tissue

On the 21st of October, 2016 animal tissue was collected post-commercial slaughter of a brahman cow from Queensland, Australia. The animal was a grey, dehorned, pure bred, registered Australian Brahman, born 2^nd^ August 2002, 14 years old. Prior to slaughter she had 12 calves. Tissue samples were collected from the abattoir, placed into labelled Eppendorf tubes and snap frozen in liquid nitrogen. Upon examination of the removed organs, it was confirmed that the cow was pregnant with an approximately 3 month old fetus, from which samples were also collected and snap frozen. Samples were transported to the University of Queensland on dry ice and stored at −80C until use.

### DNA extraction and sequencing

DNA was extracted from 71mg of kidney tissue with the Qiagen HMW MagAttract extraction kit. The total yield was 120ug of DNA as assessed by spectrometer. The HMW DNA was sent to the Ramaciotti Centre for Genomics (UNSW, Sydney, Australia) for sequencing. SMRTbell libraries were prepared using a standard procedure and sequencing on a 28 PacBio Sequel SMRTcells.

The same DNA preparation was used to generate an Illumina library using the PCR free genomic library preparation kit at the Ramaciotti Centre for genomics. The library was run across 4 lanes of a NovaSeq (Illumina), multiplexed with other samples.

Data from a MinION flowcell was obtained by preparing a 1D library from a Brahman bull, producing 4.7GB of data with a read N50 of 14.5kb. The reads were mapped to the reference using minimap2 [24] with the pre-set ONT settings.

### General computational details

Analysis was performed on the Linux Supercomputing system at the University of Queensland, Australia. Where possible, single threaded operations were parallelized using the Parallel package [25]. Statistical analysis and data visualization was performed in R [26] (version 4.0.2) in Rstudio (version 1.3.959).

### Sequence quality control

PacBio sequences were quality controlled and trimmed using the scrubbing pipeline https://dazzlerblog.wordpress.com/ which removes both low quality regions as well as missed adapters from the dataset based on pileups.

Illumina sequences were trimmed with QUADtrim [27] including removal of adapter sequences by read overlap, removal or low quality bases from both the start and end of the reads, and removal of trailing G’s, which is an artefact of the color chemistry on the current iteration of the Novaseq.

### Assembly

Contigs were assembled from the PacBio subreads using Falcon [28]. The assembly settings were: pread > 120000, daligned raw read overlap settings ‘-T16 -k14 -h35 -w7 -e.80 -l1000 -s1000 -M58 -B200’, daligner pread overlap settings ‘-T16 -k14 -h60 -w6 -e.95 -l500 - s1000 -M58 -B50’, falcon consensus calling ‘--output_multi --min_idt 0.70 --min_cov 4 -- max_n_read 200 --n_core 24’ and final overlap filtering ‘--max_diff 140 --max_cov 220 -- min_cov 5 --bestn 20 --n_core 24’.

The primary and associated contigs were combined for error correction, called polishing. The subreads bam files (which were directly obtained from the sequencing provider) were aligned to the assembled contigs using pbalign with default parameters. The alignments were then used to polish the contigs. Polishing of each contig was run on 24 cores and output included sequence files (fasta, fastq) and variant files (vfc and gff). The polished contigs were then used in a second round of polishing using the same method and parameters.

ONT data was assembled using Raven [7] and chimeric contigs were split using RagTag [8] based on comparison to the PacBio assembly of the same animal with default settings. The contigs were compared to the PacBio Assembly using Nucmer [29].

Blood from the Brahman cow was sent to Dovetail Genomics for scaffolding using Chicago and Hi-C. Dovetail Genomics HiRise [30] software was used to scaffold the assembly. Hi-C data was mapped back to the scaffolds using bwa-mem [31] for a final round of manual scaffolding. To determine which scaffolds were joined each of the independently aligned reads from each pair used. Ambiguous mapping reads were first removed from the data. The Number of expected (E) spanning reads between two contigs was calculated as

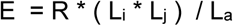

Where R is the total number of observed read pairs that mapped over 5000 bp apart, L_i_ and L_j_ are the lengths of the two contigs *i* and *j* respectively, and L_a_ is the total assembly length. If the number of observed shared links was greater than 10 times the expected number of readpairs, then the contigs were joined using a string to 1000 N’s, at the end where the density of shared links was highest. The final scaffolders were aligned to the ARD1.2 taurine genome assembly, and confirmed concordance with the genome macro-structure. Scaffolds were named based on homology to *B. taurus* [4] chromosomes.

### Gap filling

Four rounds of gap filling were undertaken with PBJelly [6]. Further rounds of gap filling did not show any improvement. Because of the possibility that short contigs were incorrectly orientated in the assembly contigs less than 1 MB in length there were flanked by gaps were reverse complemented and another 2 rounds of gap filling undertaken. Where the short contigs did not have the flanking gaps filled there were returned to their original orientation.

Data from a MinION flowcell was obtained by preparing a 1D library from a Brahman bull, producing 4.7GB of data with a read N50 of 14.5Kb. The reads were mapped to the reference using minimap2 [24] with the pre-set ONT settings. Regions with gaps were examined and three were completely covered by reads, the first was a 8.6Kb repeat on chromosome 1, the second was a 5.5kb insert, the third showed an incorrectly inserted 7.7kb region that was replaced with 320bp.

PacBio and Oxford Naopore Technologies sequences were then aligned back to the new contigs using minimap2 [24] with a reduced gap extensions penalty (-E 1,0) to allow extended gaps within reads. Alignments were filtered to only include reads of 5000 or more base pair that had less than 100bp of soft or hard clipped at either end. Where there were reads spanning the gaps the sequence was filled and the new, reassembled contig replaced the join site of the scaffolds.

### Removal of spurious indels

Illumina reads were aligned to the assembled genome using bwa-mem [31]. The alignment was then used to call variants with the mpileup command. The output file was then filtered to identify regions which showed two deletions within 5bp of each other, with a minimum total depth of 8 reads, a minimum number of reads with the deletion of 4, and ratio of between 0.2 to 0.8 of the reference (i.e. the base in the reference compared to the number of bases with the deletion). The second base position that matched these criteria for each region was removed from the contig – i.e. the reference sequence was changed from including the insertion of the indel, to the deletion. The reads were then realigned to the edited scaffold and the edited position was inspected for the retention of an indel call. The positions which exhibited insertions or deletions in at least 2 reads were manually inspected (N=52 of 583675 edited sites). Of these 27 were assessed as unimproved compared to the original sequence and the base deletion was reversed such that the original sequence was reinstated.

### GC content

The genome was first split into segments using samtools [32], which were then analysed using the geecee commands from the EMBOSS package [9] was used to calculated the GC content of each 1MB window.

### Structural variant detection

The scrubbed PacBio reads were aligned back to the reference assembly using minimap2 [24] with settings ‘-ax map-pb -A4 -B6 -O6,26 -E1,0’ to favour mapping across insertions and deletions instead of soft clipping. The resulting sam file was then sorted and indexed with samtools [32] version 1.9. SVIM [33] version 1.4 was then used to call structural variants between 10 and 1000000bp. Variants with quality scores (as determined by the software -roughly analogous to number of supporting reads) below 10 were then filtered out. As the reads were from the same animal that the reference was built, all identified structural variants therefore must be heterozygous in the animal, therefore only structural variants that were called as heterozygous by SVIM were included in downstream analysis.

### Gene placement

The nucleotide sequence of genes annotated in the EMBL ARS-UCD1.2 (GCA_002263795.2) annotation file of the B. taurus genome [4] were extracted using samtools [32]. The nucleotide sequences were then aligned to the Brahman genome using BLASTn with a minimum e-value cutoff of 10^−10^. Results were filtered with a minimum and maximum alignment length of 95% and 105% the *B. taurus* gene length, and a minimum homology of 97%. Where a gene consisted of a single loci which met these criteria it was considered to be uniquely mapped to the genome, genes with more than one loci meeting these criteria were considered to not be uniquely mapped to the genome.

### Gene clusters

To assess the probability of the observed number of genes appearing in a 1MB window by chance a random permutation was applied. The distribution of the expected number of genes within a 1MB window was calculated by randomly sampling N_*i*_ positions of genes from L_*i*_ total positions for each chromosome C_*i*_. The random sampling was performed 10,000 times for each chromosome; where N is the number of observed genes on the *i*-th chromosome, and L is the length of the *i*-th chromosome. These distributions were then used to calculate the percentiles of the number of genes in each 1MB window under the null hypothesis that the genes are randomly distributed.

### GO term analysis

The gene sequences that were placed within the Brahman genome were compared to the uniprot database [34] using BLASTx version 2.9.0 was used to align the gene sequence to the Brahman genome with a minimum e-value cutoff of 10^−10^. GO terms were extracted using BLAST2GO [35]. GO term enrichment was calculated using topGO [36] in R. GO terms with less than 5 genes were not included in the analysis. P-values were calculated using the weight01 algorithm, which includes an internal multiple testing correction.

Functional clustering of GO terms identified in the expanded and contracted gene copy number analysis was calculated using NaviGO {navigo}. As a control, a random subset of *m* genes were selected from the genome annotations, where *m* was the number of genes in the expanded and contracted gene lists. This was repeated 5 times. These genes were then input into topGO and the resulting significant GO terms (N=20 to N=28) were analysed in naviGO {Wei, 2017 #192}. With the edges calculated using Relevance Semantic Similarity (RSS) with a cutoff of 0.6, the largest cluster had 6 nodes, all others had 3 or less. This was considered the background level of connections. Functional clustering of the GO terms identified in the expanded and contracted gene sets was then identified using NaviGO with the RSS connection algorithm and 0.6 as the cut-off value. GO term definitions were obtained from EBI QuickGO{Binns, 2009 #193}.

## Acknowledgments

The authors thank the Elrose Brahman stud for providing biological samples from the carcass of the animal used to build the reference genome, and the industry contributors for the semen samples. We also thank Dr Russel Lyons for help with sample handling. The authors are grateful to Meat and Livestock Australia for providing the funding for this study under projects (P.PSH.0868 and L.GEN.1808).

